# 3D X-ray microscopy lights up nanoparticles in plants

**DOI:** 10.1101/2025.05.08.652794

**Authors:** Emil V. Kristensen, Francesca Siracusa, Andrea Pinna, Idil Ertem, Augusta Szameitat, Julie Villanova, Xiaoyu Gao, Adrian Schiefler, Dominique J. Tobler, Subhasis Ghoshal, Søren Husted, Rajmund Mokso

**Affiliations:** Technical University of Denmark, Department of Physics, Fysikvej, building 307, 2800 Kgs. Lyngby, Denmark; University of Copenhagen, Plant and Environmental Sciences, Thorvaldsensvej 40, 1871 Frederiksberg; ESRF - The European synchrotron, 71 Avenue des Martyrs, 38000 Grenoble, France; McGill University, Department of Civil Engineering, 817 Sherbrooke Street West, Montreal, Quebec H3A 0C3, Canada; Baylor University, Department of Environmental Science, One Bear Place 97266, Waco, TX 76798–7266, USA

## Abstract

The discovery of novel plant fertilization strategies heavily relies on our capabilities to probe physiological processes in living plants with sub-cellular precision. State-of-the-art microscopy techniques are in general limited to surface investigation or they require elaborated tissue preparation and often destruction. X-ray microscopy has the potential to resolve some of these limitations by generating micro-to nanometer-scale 3D images deep into the tissue. We introduce experimental designs and the quantitative analysis methodologies, pioneering nanoscale (≈150 nm resolution) in-vivo 3D microscopy of plant tissue. We show the first direct *in-vivo* visualization of foliar-applied untagged nanoparticulate fertilizers deep under the leaf surface, not accessible by other microscopy methods. Ultimately, our approach provides the means for a direct observation of nanoparticle transport and dissolution in living plant tissue, a step critical for developing sustainable plant fertilization approaches.

## Introduction

A major issue in agriculture is the inefficiency of traditional fertilization strategies due to immobilization in soil and leaching to aquatic environments causing negative impacts to the local environment.^1^ Foliar application of the fertilizer, i.e. direct application of the fertilizers to the leaf, has the potential to bypass the limitations of the soil system while providing readily available nutrients to plants and reducing nutrient losses to the environment. However, leaf scorching induced by the application of soluble salts, and in some cases, the poor re-mobilization of foliar applied nutrients, often reduce the efficiency of the foliar fertilization practices.^2^ Introducing nanotechnology to the field of plant science opens up entirely new ways of delivering nutrients to plants. Due to their small size and customizable surface properties, nanoparticulate (NP) fertilizers can deliver essential plant nutrients in ways that differ significantly from raw mineral ions. This method can help bypass the limitations of conventional foliar fertilization strategies ^3,4^. Furthermore, NP physicochemical properties can be tuned to allow for improved leaf uptake and translocation, and more advanced functionalization can be introduced to enable e.g. stimulus-responsive release of nutrient cargo allowing for slow and programmed release. While studies show the uptake and beneficial effects of foliar application of NP fertilizer^5,6^, very few of these studies truly show how the NPs pass major transport barriers, such as crossing the leaf surface, entering the phloem, and translocating through to other plant parts. ^7–9^ Imaging pristine and untagged NP clusters at the sub-cellar resolution level is essential to understanding how particles are taken up and transported within the plant structure. The development of techniques able to image native NPs and their transport and aggregation/dissolution behavior within plant tissue will allow to clarify some of the barriers in designing efficient NFs ^10,11^.

One of the emerging techniques for plant microstructure characterization is X-ray computed micro-tomography (*µ*CT)^12–18^. Common investigation areas for X-ray CT are the characterization of vascular tissue and the transport phenomena in both phloem and xylem ^19–21^. Other typical studies involve root growth or *in-vivo* high throughput systems where the same plant is scanned many times tracking developments as the plant grows ^22,23^ and, investigations of seed structures and growth. ^24^ The uptake of NPs through the root system has been investigated by the use of nano-tomography techniques ^25–27^. These results fall into two categories: either micro-CT is used to image living tissue structures at the cellular level, without the ability to resolve sub-cellular structures, or nanoscale resolution is achieved, but only for fixed tissue samples.

When working with living organisms the resolution limit is not only given by the instrument design but also by the physical effect of the ionizing radiation, because radiation dose scales with the spatial resolution as a power of four. Therefore, nanoscale X-ray imaging typically causes the destruction of the living tissue. For this reason, nanoscale-resolution studies do not only require technically advanced X-ray setups but equally well optimized sample preparation procedures. The highest achievable 3D resolution using X-rays is obtained through ptychography at synchrotrons, ^28^ however, this technique requires sub-millimeter sample sizes and acquisition times of several hours. Alternatively, scanning electron microscopy (SEM) is also widely used to achieve sub-micrometer resolution imaging of plants. ^29^ But due to SEM being a surface probing technique, investigating internal structure requires sectioning of the sample (ie FIB-SEM), making it a destructive technique. Owing to its unparalleled resolving power, transmission electron microscopy (TEM) can be used to image internal features of cells at the nanoscale level, though extensive/tedious sample preparation procedures and the requirement of ultra-thin dehydrated sections make this technique unsuitable for mapping dynamic processes occurring in plant tissues. ^30^

Easy procedures for sample preparation and cellular resolution make confocal laser scanning microscopy (CLSM) a powerful technique for *in-vivo* imaging of plant tissues. However, the imaging depth of CLSM is limited to the superficial layer of the sample. Scattering, photobleaching, and noise are a few factors that restrict its use to study processes occurring a few hundred microns below the leaf surface while also requiring fluorophores, whether naturally present in the sample (e.g., chloroplasts), or conjugation with a reporting tag, such as a fluorophore or Raman probe^23,31^. Since X-ray imaging is based on local density differences within the sample, it is often limited by the uniform and low-density nature of plant tissue. ^32^ Often this is mitigated by elaborate sample fixation and staining using contrast agents, which in turn alter the sample to some degree. ^33^ In this study, we avoided the use of contrast agents and, whenever possible, conducted our work without tissue fixation. Instead, we achieve high definition 3D images by fully exploiting the coherence properties of modern synchrotron X-ray sources, enabling phase contrast imaging to resolve small density differences between various tissue types in plants.

We present three synchrotron X-ray phase contrast tomographic microscopy methods for imaging plant tissue and NPs behavior inside plant tissue (illustrated in Fig 1). In rather large (≥5×5 mm^2^) freeze-dried leaf samples we were able to zoom in to sub-cellular structures as small as thylakoid stacks inside chloroplasts deploying the method of lens-based Transmission X-ray microscopy (TXM)^34^ with near 100 nm isotropic spatial resolution in 3D. Larger regions (≈ 1 cm^2^) of living plant leaves were scanned using parallel beam *µ*CT with ≈ 1*µ*m m isotropic spatial resolution. Subcellular structures and NP aggregation *in-planta* in living plants were captured for the first time with nano-holotomography.^35^ Applying these modalities, we gained insights into how NPs are taken up and clustered after foliar application on plant leaves.

**Figure 1:**
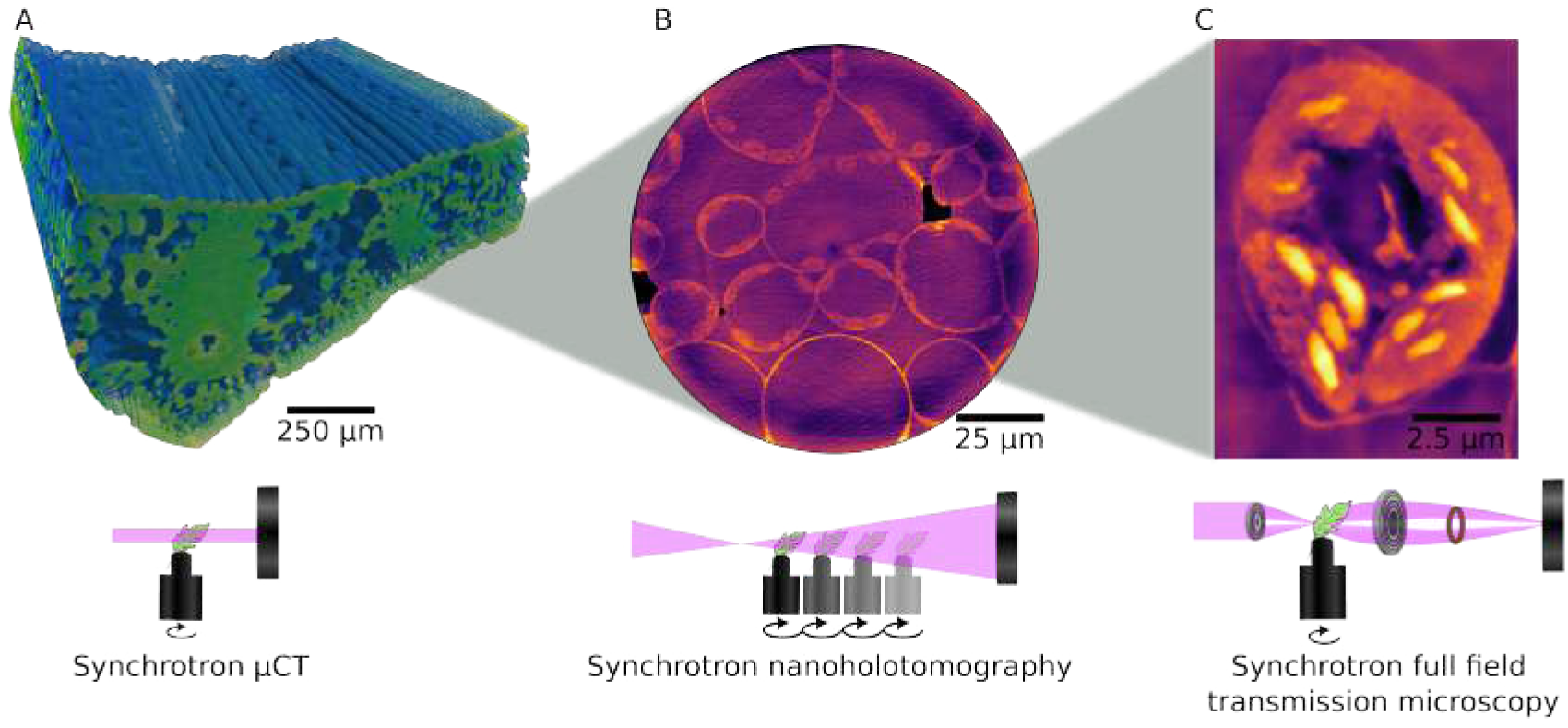
Schematic illustrating the three synchrotron tomography techniques utilized and their corresponding results. **A** Parallel beam *µ*CT, with a 3D reconstruction of a barley leaf. **B** Nano-holotomography, featuring a virtual slice from a CT scan of a living barley leaf, revealing sub-cellular structures. **C** Full-field X-ray transmission tomography, with a slice depicting a single cell with sub-cellular organelles visible.

## Results

By optimizing and applying three phase contrast synchrotron X-ray microscopy methods on freeze-dried and living leaf tissue we obtained 3D volumetric representation of barley, soy and tomato leaves from *µ*m to tens of nm voxel sizes. Utilizing phase contrast methods significantly increases dose efficiency compared to absorption-based techniques ^36,37^. Using these methods, we describe anatomical structures ranging from leaf pore spaces in the entire plant organ down to subcellular organelles such as thylakoid stacks inside chloroplasts, as wells as NP aggregation inside living leaf tissue.

### 3D X-ray nano-imaging of dehydrated plant tissue

A critical step in high-resolution 3D imaging is to assure the stability of the sample at the level of the desired resolution during the scanning process. For nanoscale imaging, such as electron microscopy, the plant tissue is typically fixed and/or freeze-dried. The latter assures that there is no water in the sample to compromise the stability during exposure to the probing radiation/particles. Freeze-dried soybean leaves were imaged with TXM to demonstrate the nanoscale 3D visualization within samples several orders of magnitude larger than what is achievable with electron microscopy. The experiment was carried out at the BL47XU beamline at the SPRING8 synchrotron. First, the sample was imaged with synchrotron *µ*CT with 385 nm pixel size to create overview images of the entire freeze-dried leaf with a field of view of 1×2 mm (Fig. 2 **A-B**). The reconstructed data from the *µ*CT images enabled the identification of a suitable location for the high-resolution scans. These images contain structural information about the plant tissue and the organization of cell types in 3D, despite being at a low resolution not allowing for visualization of sub-cellular structures. Epidermal cells at the surface, palisade mesophylls, spongy mesophylls, trichomes, and vascular tissue with both xylem and phloem can be identified (Fig. 2 **A-B**). Selecting the palisade mesophyll cells at the location indicated by the circle (Fig. 2A), the TXM instrument was used with a 50 nm pixel size to obtain sub-cellular details of the marked region (Fig. 2C)^1^.

**Figure 2:**
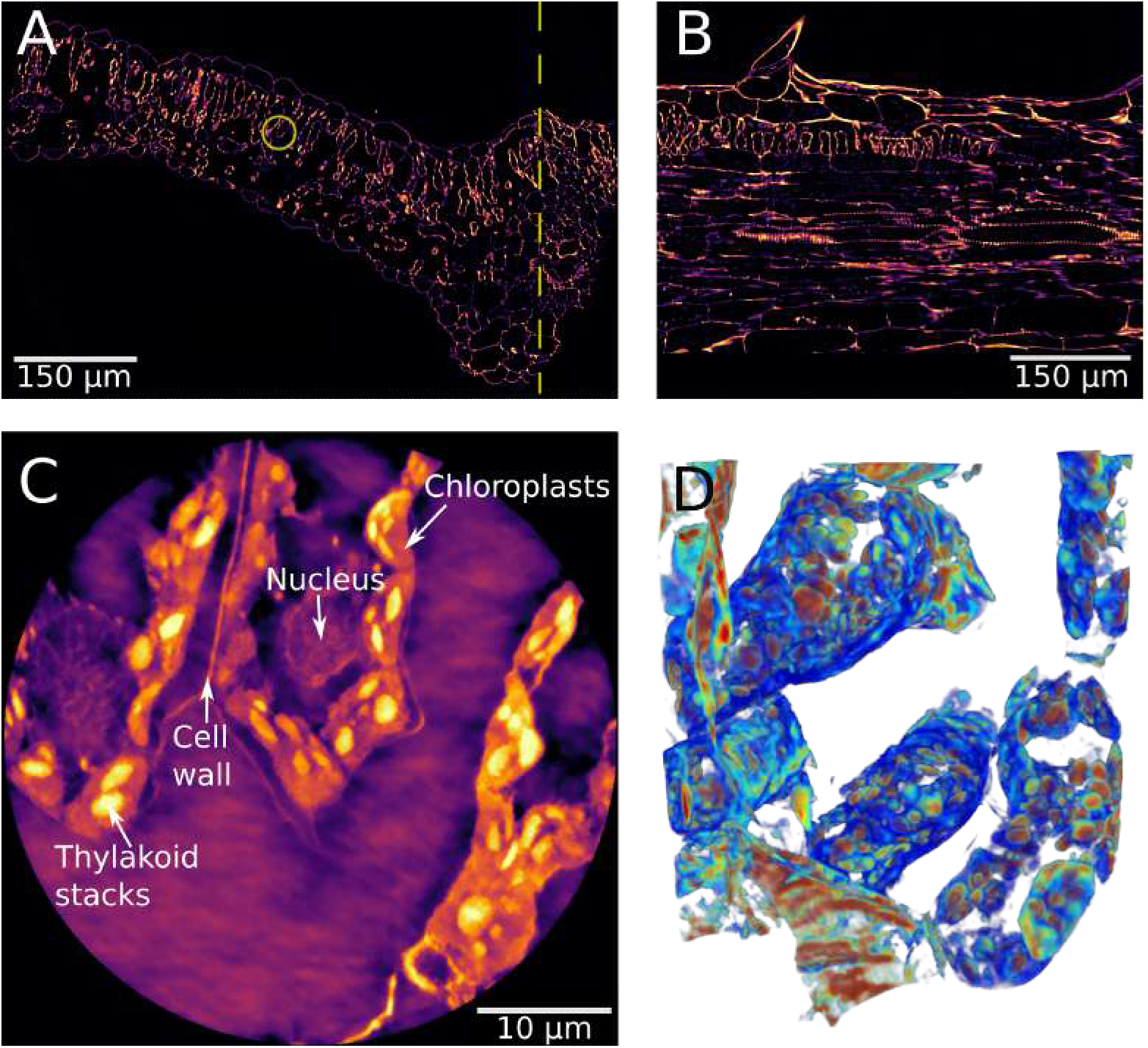
Synchrotron *µ*CT and TXM scans of freeze-dried soybean leaf. **A)** Low resolution (0.385 *µ*m pixel size). The dashed line indicates the position of the orthoslice visible in **B** and the circle indicates the high resolution scan in **C-D**. **B)** Orthoslice of the same scan as **A**, cross-section of vascular tissue and trichome visible. **C)** Nano-resolution image of the region marked in **A** with subcellular organelles labeled. **D)** 3D rendering of the nano-resolution scan in **C)**. Movies of 3D rendering are available in S2-S3.

Three elongated palisade mesophyll cells are seen with clearly visible sub-cellular structures. A 3D rendering of these cells (Fig 2**D**) was made to highlight the 3D organization of tissues, such as the nucleus, cell walls, chloroplast membranes, and the stroma, in green to blue colors. Following this, the denser thylakoid stacks inside the chloroplasts were colored red.

Surrounding the cells, a thin line representing the cell wall is visible. In each cell, distinguishable chloroplasts are visible as medium-contrast regions near the edge of the cell filled with the highest-intensity disk-shaped spots, identified as thylakoid stacks. Inside the cells, the nucleus can be seen. Generally, the tissue is observed to be almost entirely intact except for the slight distortion of cell shape due to dehydration from sample preparation.

### Tomographic imaging of hydrated plant organs

The first step in our study is the mapping of large volumes of the plant leaf at the micrometer scale. For this we used synchrotron *µ*CT at the DANMAX beamline of the MAX IV Laboratory. The leaf of an intact, living barley plant was imaged in 3D with an isotropic voxel size of 550 nm, and the measured spatial resolution of ≈ 1.5µm (Fig. 3). These images reveal the cellular organization of the leaf including mesophyll cells, epidermis, vascular bundles with xylem and phloem, and stomata in the epidermis of the leaf. Although conventional *µ*CT is a well-established and widely used technique in plant biology, it still holds significant potential as a tool for further discoveries,^38^ especially when considering that such micrometer scale 3D maps of 2×2×2 mm region are acquired in less than a minute.

**Figure 3:**
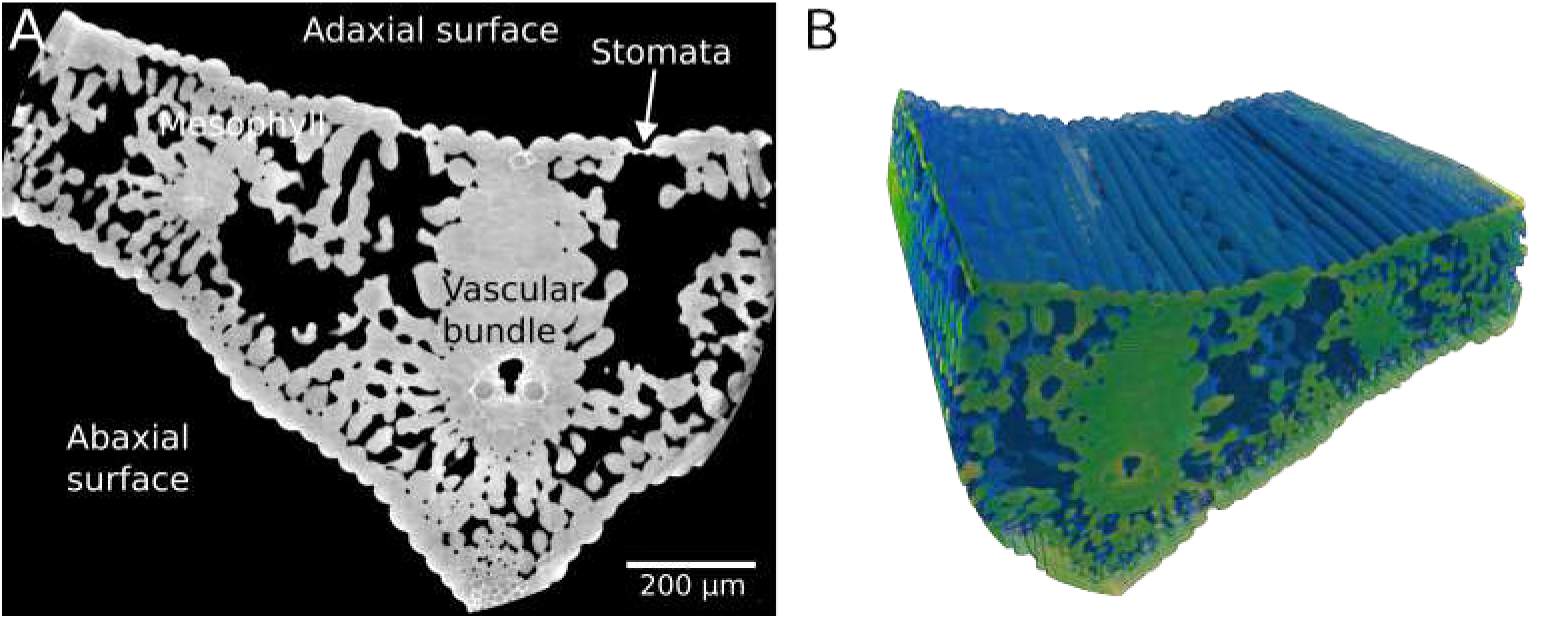
Synchrotron *µ*CT images of barley leaf from the DANMAX beamline of MAX IV. **A** Virtual slice of barley leaf. Visible structures include stomates, mesophyll cells and vascular tissue cells xylem and phloem. **B** 3D rendering of barley leaf tomogram. Movies of 3D rendering available in S4.

In contrast to the parallel beam geometry of synchrotron *µ*CT Fig. 1A, a cone beam setup in projection geometry, as shown in Fig. 1B, enables multiscale imaging by continuous zooming in and out on the sample. ^35^ Here, the nano-holotomography approach was used to gain an order of magnitude in resolution, yet keeping the plant alive during image acquisition. In the following, we utilized the nano-holotomography setup of the ID16B beamline at ESRF. Living barley leaves were visualized, first untreated (Fig. 4A**-B**) where we can identify internal cells in bright and air-filled pore space in dark gray values. Despite the high resolution (100 nm pixel size), no sub-cellular structures (chloroplasts, nucleus, cell walls, etc.) are visible. However, we made those visible by infiltrating the barley leaf with Milli-Q water which replaces the air inside the leaf, and matching the density of the cell contents. In particular, denser structures of chloroplasts and cell walls become visible (Fig. 4**C-D**). Fig. 4**E-F** highlights the vascular tissue with xylem and phloem cells where the main contrast is seen due to the density differences between the heavy vascular cell walls and the cells content. Similar phase matching effects were previously observed in grating-based ray phase-contrast imaging, where biological samples are embedded in paraffin to reduce the too large X-ray phase shift variation between the tissue of interest and the surrounding medium ^39,40^.

**Figure 4:**
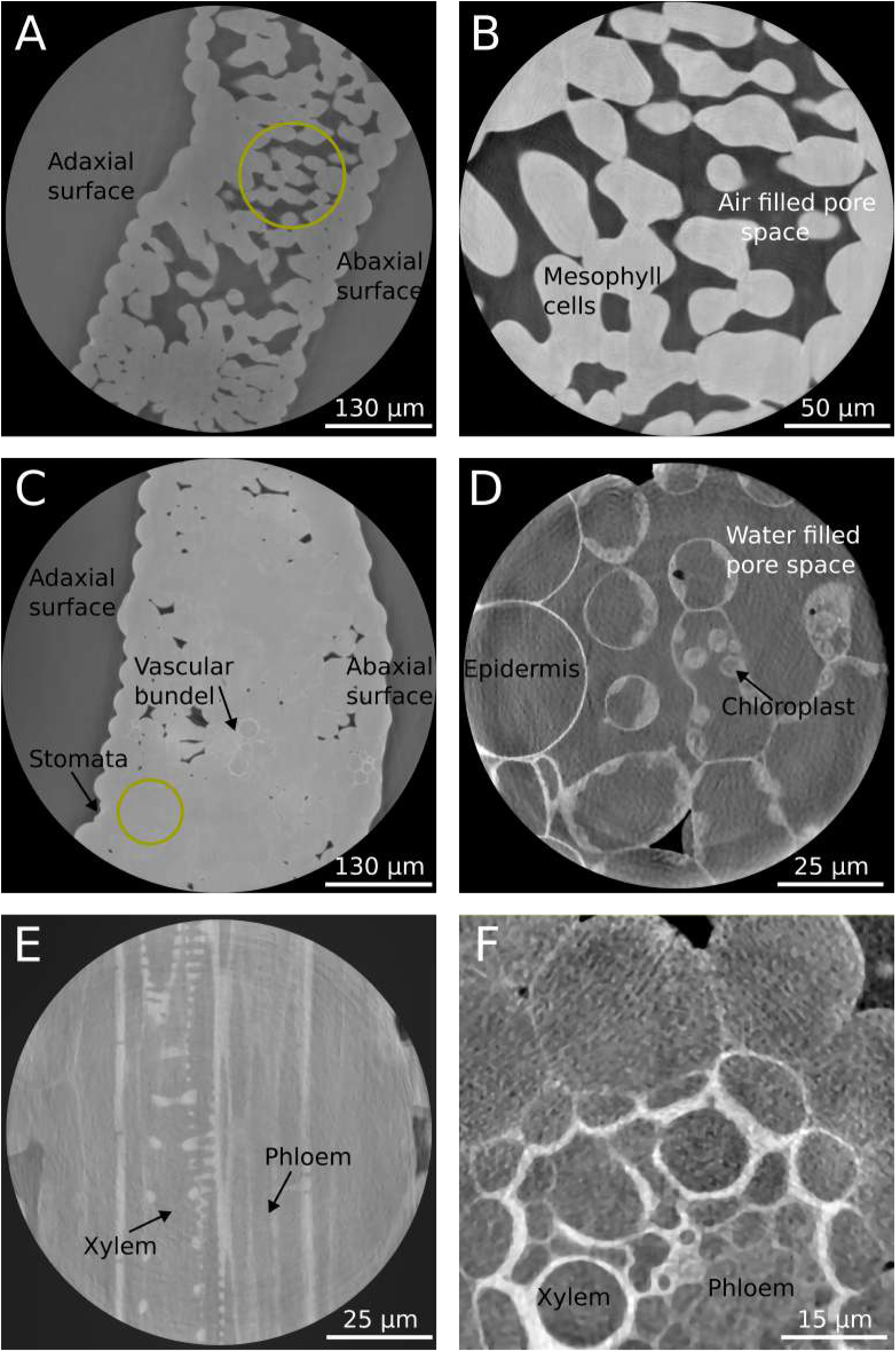
Synchrotron nanoholotomography images of living barley leaves. **A** Low-resolution scan of an untreated leaf sample (257 nm pixel size). **B** High resolution scan of the mesophyll region indicated in A (100 nm pixel size). **C** Low-resolution scan of water-filled Barley leaf (257 nm pixel size). **D** High-resolution image of the water-filled mesophyll tissue marked in C (50 nm pixel size). Chloroplasts and cell walls clearly visible. **E** and **F** Orthoslices of high-resolution images of midvein tissue with clearly visible Xylem and Phloem cells (50 nm pixel size). Movies of 3D rendering available in S5-S8

Next we investigated the aggregation behavior of manganese oxide (MnO) NPs coated with polyacrylic acid (nPAA-MnO) in barley leaves. nPAA-MnO (characterized in SI Fig S1) have previously been used as model Mn containing nano-fertilizer due to their proven ability to be taken up by leaves and provide bioavailable Mn to the plant, triggered by a drop in pH once entering the mesophyll apoplast. ^41^ The uptake of DiI fluorophore-labelled nPAA-MnO (DiI-nPAA-MnO) 5 hours after drop deposition in barley leaves has been demonstrated using CLSM (See SI Movie S1). In our experiment, the MnO suspension was infiltrated into the leaf using a syringe to ensure uptake and then scanned within 4 hours after application. Chloroplasts and cell walls are visible in this sample (Fig. 5A), but there are no discrete NPs visible inside the plant tissue. The individual NPs are smaller (≈ 20 nm particles) than the spatial resolution of the setup, therefore if the particles are dispersed and do not aggregate, we can not visualize them. This observation is consistent for all replicates of this experiment type, regardless of whether the particles were infiltrated into the leaf, or applied as droplets on the surface. To validate this hypothesis, we forced the MnO NPs to aggregate and form clusters larger than the spatial resolution of the setup (i.e. cluster size *>*150 nm). A barley leaf was first infiltrated with CaCl_2_ solution, followed by MnO NP solution infiltration. Here the Ca^2+^ ions act as flocculant. This approach was successful as shown in Fig. 5**A** and **B**, where we now see 0.5-10 *µ*m sized bright spots located within, or just next to, the cell wall. To verify that these aggregates are indeed the dosed MnO NPs a control only infiltrated with the CaCl_2_ solution was also scanned. Notably, no bright spots are seen in leaves treated with CaCl_2_ solution or water only, indicating that the bright spots are MnO NP clusters (Movie S12).

**Figure 5:**
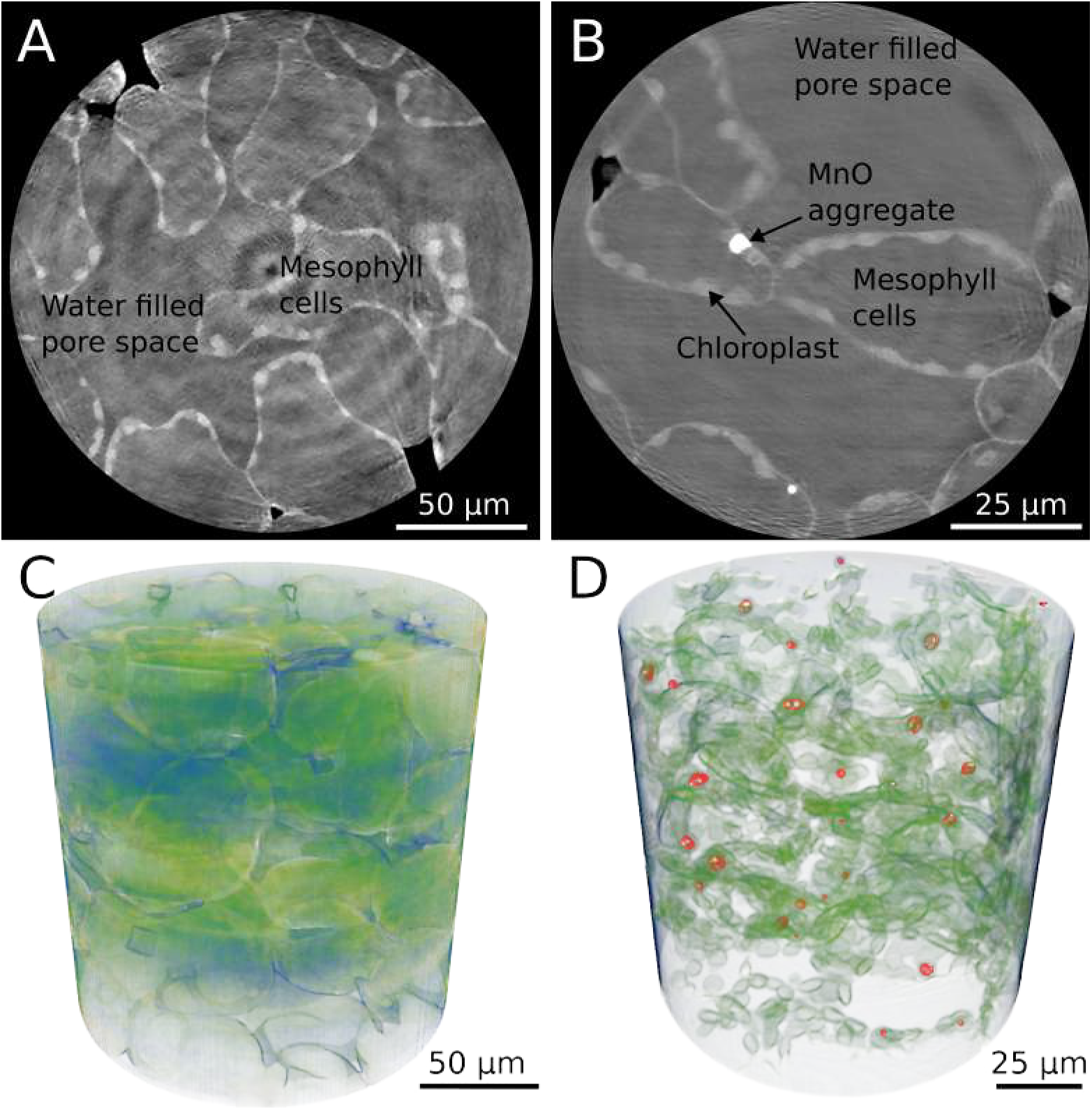
Synchrotron nano-holotomography scans (100 and 50 nm pixel size) of barley leaves infiltrated with MnO NPs. **A** Mesophyll cells of MnO infiltrated leaf. No particle clusters are visible. **B** Mesophyll cells of barley leaf infiltrated with CaCl_2_ before MnO infiltration. Particle clusters are visible inside cell walls. **C** and **D** 3D visualization of **A** and **B** with MnO particle clusters shown in red. Movies of 3D rendering available in Movies S9-S10

Besides using bare NPs as micronutrient fertilizers, nanomaterials can also be encapsulated in other carriers for a controlled release function and higher uptake efficiency. For example, the transport efficiency of ZnO NPs has been previously shown to be enhanced when encapsulated in a mesoporous silica nanoshell (MSN)^9,42^. These MSN-coated particles are denser, larger (≈ 70 nm diameter), and tend to aggregate more than the uncoated MnO NPs. For these reasons, the MSN particles are more visible and easier to image.

A liquid suspension (containing 0.02% Silwet Gold for lowering surface tension) of ZnO-filled MSN particles was applied in drops to the surface of a tomato leaf. CT images of tomato leaves were taken three hours after the application of MSN droplets to the leaf surface, under the same imaging conditions as the barley experiments, revealing aggregation at the application site (Fig. 6). Additionally, aggregates were observed within the cell walls of the leaf.

**Figure 6:**
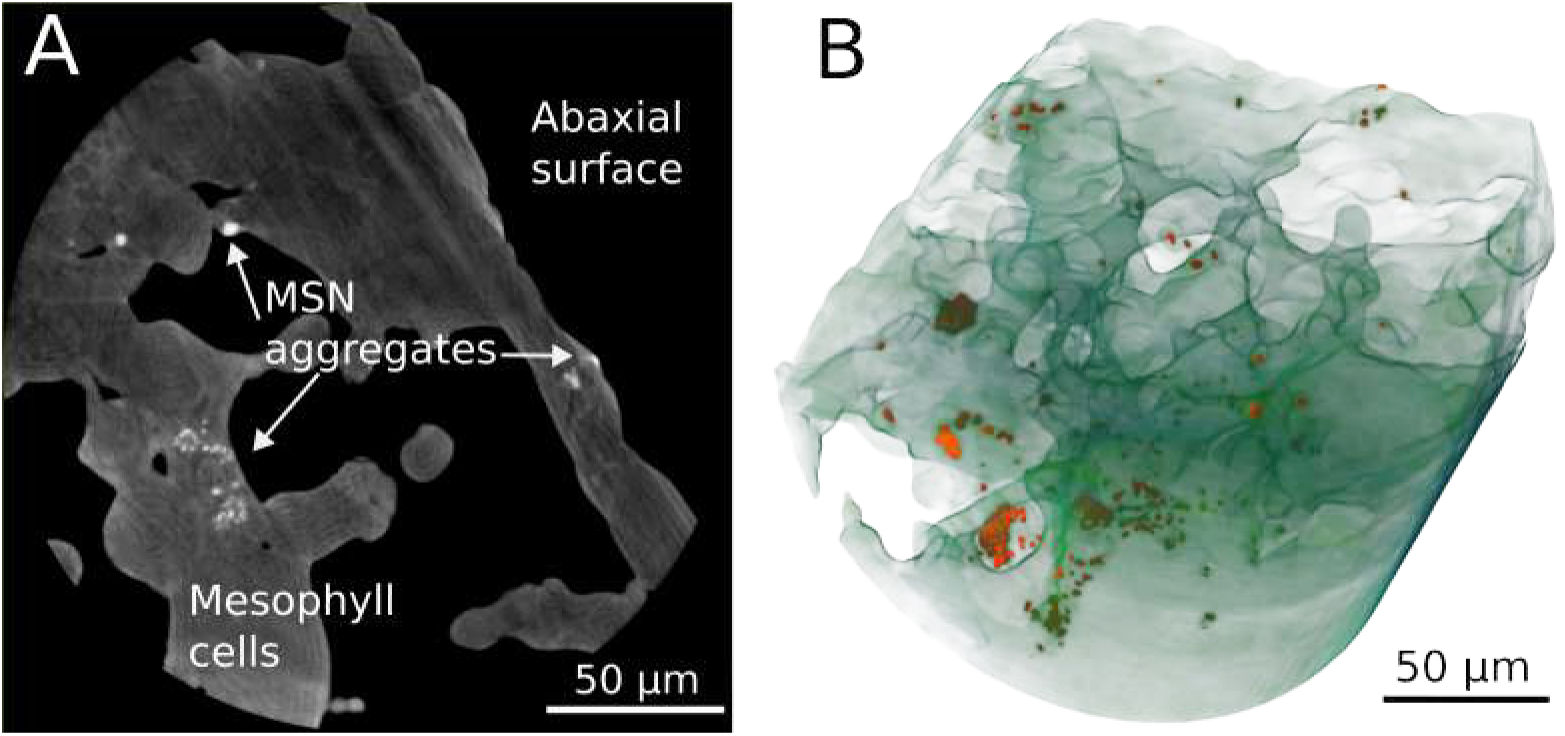
Synchrotron nano-holotomography scan (100 nm pixel size) of tomato leaf dosed with ZnO filled MSN particles. **A**) Virtual slice showing MSN particles aggregated in the apoplast as bright spots. **B**) 3D visualization of NP clusters (red) with leaf tissue in green. Movies of 3D rendering are available in Movies S11.

### Quantitative analysis

The theoretical resolution of the presented microscopy techniques is mainly given by the optical design and acquisition method, while the actual image quality is often limited by the mechanical stability of the sample and the maximum acceptable radiation dose. In addition, the contrast ratio between tissue types in the sample can also affect image quality. Both for the TXM and nano-holotomography methods, the smallest pixel size achieved is 50 nm. In studies of hard materials with ample contrast, this resolution can often be fully utilized revealing features at the smallest resolvable size^43,44^. For living biological tissues, this is more difficult due to the inherent movement of living organisms, especially when exposed to radiation.^45^ As seen in figure 4B, the high resolution is not necessarily enough if the features of interest cannot be imaged at sufficient contrast to distinguish them. This leads us to evaluate the resolution using a Fourier-based correlation method ^46^ (Tab. 1). As expected, the *µ*CT (Fig 3) scan results in a ≈ 2*µ*m resolution. For the nano-imaging methods, we measured the resolution of ≈ 149 nm for the TXM compared to ≈ 168 nm using the nano-holotomography setup. While the effective voxel size is the same (50 nm) the small difference in actual resolution is due to instabilities expected of the living sample during scanning in nano-holotomography, in contrast to the freeze dried samples used in the TXM measurements. These values are similar, but the main qualitative difference between the scan of dehydrated and hydrated plants is the available contrast. In the absence of water, the tissue-to-air contrast in the TXM images (Fig. 2) allows for a simple threshold segmentation of individual organelles. The water and cell body electron density is very similar and therefore pose a major challenge for label-free imaging methods, such as those used in this study. Thus the image of living tissue with holotomography shows less contrast between organelles and the rest of the cell.

**Table 1:** The measured resolution and pixel size for each imaging method.

| Image | Pixel size nm | Resolution [nm] |
| --- | --- | --- |
| μCT (Figure 3) | 550 | $2082 \pm 307$ |
| TXM (Figure 2) | 50 | $149 \pm 6.5$ |
| Holotomography (Figure 5B) | 50 | $168 \pm 19.2$ |

By exploiting the quantitative nature of X-ray nano-holotomography, together with the known densities of water and air, the actual electron densities of all the visible plant compartments can be measured. To do this the measured intensities of all scans containing both water and air regions were normalized as

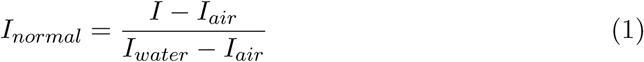

This normalization makes the gray values an indicator for local density with *ρ_air_* ≈ 0, *ρ_water_* = 1, and the denser tissue types and particles with *ρ >* 1 creating a grayscale qualitatively similar to the Hounsfield scale used in medical imaging to quantify density variations ^47,48^. The only requirement for this quantitative normalization is that both air and water is present and segmentable.

Histograms of normalized pixel values for two nano-holotomography scans are given (Fig. 7 C). The different types of tissue can be identified by the peaks in the histogram. The red curve represents the barley leaf dosed with CaCl_2_ and MnO NPs (Fig. 5B) where both the chloroplast and the cell walls can be seen as a shoulder of the water peak at gray values just above 1. At intensities of ≈ 1.6 times the water level a clear peak of the MnO NPs can be seen. The blue line represents the histogram for the scan of the barley vascular tissue (Fig. 4 **E**) with a clear shoulder from the higher densities of the vascular tissue around 1.4 times the water gray level.

**Figure 7:**
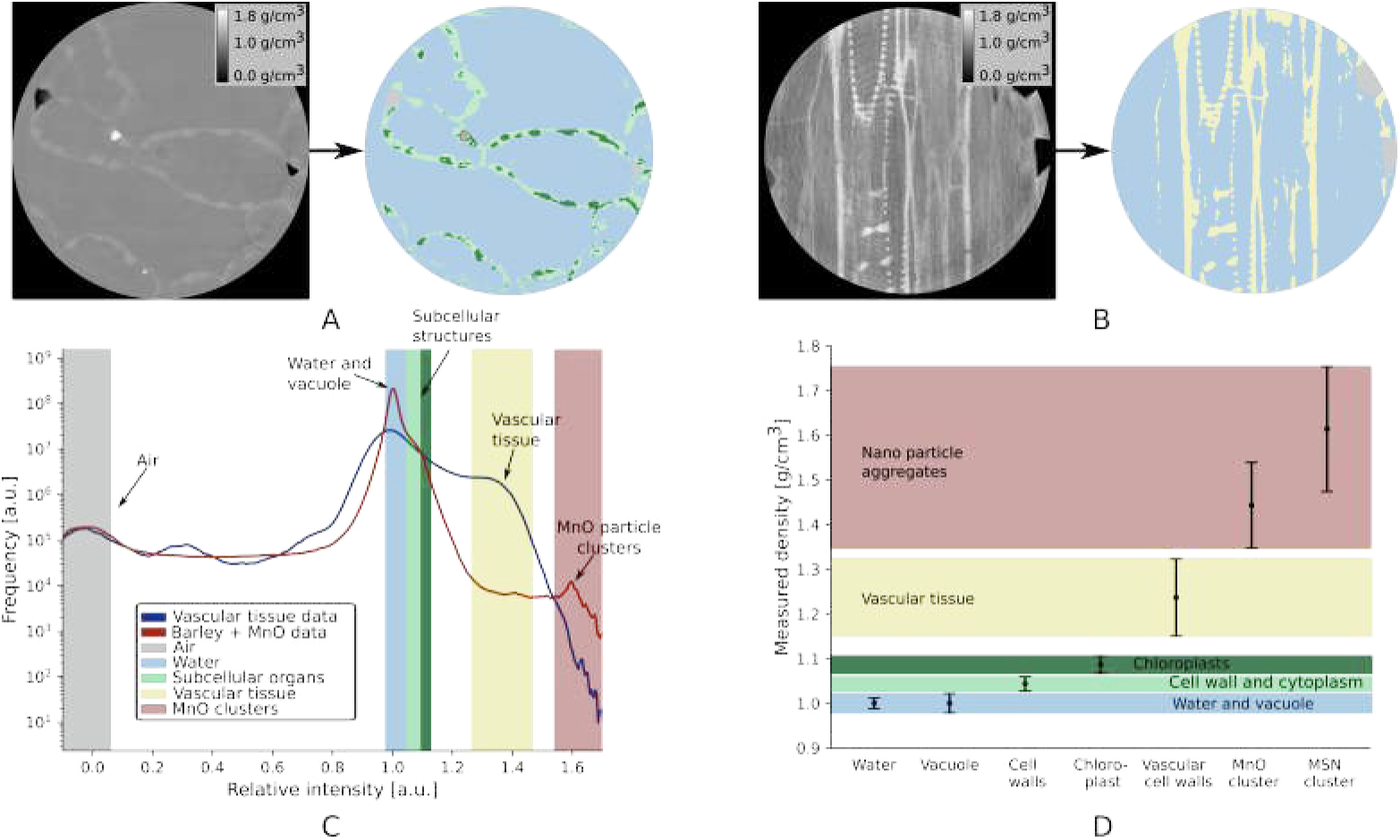
Quantification of local densities from image grayscale. **A,B** Slice of scan (Fig 5**B** and 4 **E**) along with segmented tissues in color. **C** Histogram of the pixel values of the scans (Fig. 5**B** and 4 **E**). Intensities are normalized to the mean gray level of the water. Approximate values of the different tissue types are indicated with the color bands at the value of the peak/shoulder originating from the tissue type. **D** Measured densities of different phases in the nano-holotomography scans. Water and vacuole with densities close to 1 g/cm^3^ marked with the blue area. Subcellular structures (mesophyll cell walls and chloroplasts) are marked with the green/dark green regions at densities of 1.02-1.12 g/cm^3^. Cell walls of the vascular tissue are marked with yellow at densities of 1.14 to 1.32 g/cm^3^. The MnO and MSN clusters are marked in red at the highest densities with densities of 1.35 to 1.75 g/cm^3^. Error bars indicate the standard variations in gray levels of each tissue type.

From this histogram, we can see that air and NP material can be segmented by simple thresholding as the gray values are separated enough from the rest of the tissue to avoid overlap. For the different tissue types, the gray values overlap with the values found in the water phase between the cells and inside the vacuole. For these tissue types, the segmentation is done by a random walker segmentation method. ^49^ The segmented areas are then quantified by their mean pixel intensity, translated into density (Fig. 7 **D**). The densities of the water phase (outside of cells) and vacuole overlap entirely and cannot be distinguished from these scans. At slightly higher densities chloroplasts and mesophyll cell walls are found with calculated densities of 1.08 and 1.04 g/cm^3^ respectively. At 1.25 g/cm^3^ the vascular cell walls are the densest naturally occurring tissue in the barley leaf and finally the MnO and MSN NP aggregates are measured at 1.44 and 1.61 g/cm^3^, respectively. The difference between measured density and the true density of the particles can be explained by a porous packing of the aggregates, which reduces the density.

## Methods

### Plant growth

Plants were germinated in vermiculite afterwards seedlings were transplanted to aerated 5L hydroponic containers supplied with nutrient solution containing KH_2_PO_4_ (200 *µ*M), K_2_SO_4_ (200 *µ*M), MgSO_4_·7H_2_O (300 *µ*M), NaCl (100 *µ*M), Mg(NO3)_2_·6H_2_O (300 *µ*M), Ca(NO_3_)_2_·4H_2_O (900 *µ*M), KNO_3_ (600 *µ*M), Fe(III)– EDTA–Na (50 *µ*M), H_3_BO_3_ (2 *µ*M), Na_2_MoO_4_·2H_2_O (0.8 *µ*M), ZnCl_2_ (0.7 *µ*M), MnCl_2_·4H_2_O (1 *µ*M) and CuSO_4_·5H_2_O (0.8 *µ*M). Nutrient solutions in the pots were renewed weekly and pH was maintained at 5.5 ± 0.5 using HCl. Plants were kept in greenhouse conditions with minimum day/night temperatures of 18 *^◦^*C/15 *^◦^*C and a day/ night cycle of 16 h/8 h with minimum light intensity of 500 *µ*mol photons/(m^2^s) .

### Freeze-dried samples

On day 21 after transplanting, the soybean plants were harvested. Following harvest, leaf pieces were cut into 3 mm segments and immediately frozen in hexane at -80°C, cooled with dry ice. The samples were then freeze-dried (Christ Alpha 1-4, Germany) at 1 mbar and -40°C. After preparation, the samples were stored in a sealed container.

### In vivo samples

On days 7 to 10 after transplanting, tomato and barley plants were transferred to 50 mL containers filled with vermiculite moistened with the nutrient solution from the hydroponic containers. During transportation, plants were contained in a box and supplied with light. Barley leaves were infiltrated 5 hours before analysis with 20 mM CaCl_2_ following infiltration with MnO solution (details on particle formulation are available in SI). Infiltration of solutions into the leaves was performed by applying gentle pressure between the leaf and the tip of a disposable 1 mL syringe, while slowly introducing the liquid into the leaf tissue.

Tomato seeds were rinsed three times with 18.2 MΩcm ultrapure Milli-Q water before being germinated in sterile Petri dishes lined with wet filter paper. The dishes were kept in the dark at room temperature for 10 days. Seedlings that germinated uniformly were then transplanted into aerated, light-impermeable black hydroponic units and cultivated under controlled greenhouse conditions. Each hydroponic unit, which held four plants, was filled with Hoagland solution containing all essential plant nutrients (except for Zn, keeping the plants Zn deficient) and aerated using a stainless-steel medical syringe. About 1 week after germination plants were dosed with NPs and imaged. Droplet application of NP solutions containing 0.1% Silwet Gold to leaves was administered by dropping 50 *µ*L on the leaf surface using a syringe. For tomato plants, droplets were applied to the abaxial side due to previous results showing higher uptake capabilities compared to the adaxial side. For analysis, a leaf was tightly secured without damage, in a specially designed sample holder while the roots were kept moist.

### Transmission X-ray microscopy

TXM images were taken at the BL47XU beamline at SPIRNG8 at a beam energy of 15 keV. Prior to the experiment, the *µ*CT and TXM detectors were aligned to have corresponding centers ensuring that the center position of the low-resolution images corresponds to the center of the high-resolution images. Freeze-dried samples were mounted in the sample holder using super glue and a pin making sure the region of interest on the sample was just above the pin for maximum stability of the sample. The sample holder was then placed on the sample stage with the beam station in *µ*CT mode, meaning that the condenser lens, Fresnel objective lens, and phase ring were all moved out of the beam and replaced by a 2048×1504 pixel detector just behind the sample. With this setup standard parallel beam tomography achieving a pixel size of 385 nm and a field of view of 0.79×0.79×0.58 mm^3^ is possible. The sample was then aligned to the region of interest and a full 1004 projections tomography was measured. With the *µ*CT reconstructed we then chose a new sub-region of high interest for the high-resolution scan and then align the sample to this new point in all three axes of the sample stage. The *µ*CT detector was then replaced with the TXM optics and a series of tomographic scans each with 1751 projections over 180 degrees, each with a 100 ms exposure time, was then conducted. The total scan time per tomogram amounted to around 10 min. From this procedure, a set of 920×920×784 pixel tomograms with a pixel size of 50 nm and a field of view of 40 *µ*m was measured at the region of interest chosen on the low-resolution image. From the raw projection images the 3D tomographic images were reconstructed using a filtered back-projection type algorithm implemented at the B47XU beamline at SPRING8.

### Synchrotron ***µ***CT

The DANMAX (MAX IV) beamline was set to an X-ray energy of 25 keV. A 2176 X 2176 pixel Optique Peter detector with a 10x objective was used at a sample-to-detector distance of 30 mm (closest possible due to sample holder constraints). Tomographic scans were then done by collecting 2001 projections over 180 degrees with an exposure time of 50 ms per projection. All projections were then phase-retrieved using Paganin-based phase retrieval and then tomographic rreconstruction is done using Tomopy.^50^

Full size, intact and living barley plants were mounted upside down in a custom build sample holder. The leaf of interest was then secured by clamping between two screws to hold the leaf flat and still. The roots were fixed to the top of the sample holder and kept moist by wrapping them in wet tissue paper. The walls of the sample holder consist of a Kapton tube making the sample holder transparent to X-rays from 360 degrees.

### Nano-holotomography

The nano-holotomography experiment at ESRF id16b was done at an X-ray energy of 29.9 keV with the beam focused by a set of KB mirrors to a focus spot size of 50 nm. Sample leaves were prepared by cutting ≈ 10 mm x 50 mm slice of the leaf which was then placed in a sealed pipette tip with a small amount of water to keep the leaf section alive. The holotomography scans were then done by collecting tomograms at 4 different focus to sample distances and 900-2505 projections over 360 degrees (varying from scans) using a PCO edge detector with 2048×2048 pixels. At the highest resolution of 50 nm pixel size the FOV were around 100 *µ*m. Typical scan times for a four-distance tomogram were around 40 min to 1 hour. All projections were then resampled to the lowest pixel size and phase retrieved using Multi-distance Paganin’s method ^51,52^ with *δ/β* = 85 and iterative refinement. After this, tomographic reeconstruction using PyHST2^53^ results then in the final 3D datasets.

### Post processing and visualization

The tomographic data was then post-processed using a ring-removing algorithm, the manual 3D-RBF function from Dragonfly to correct intensity gradients and finally, the tomograms were denoised using a combination of median filter and an iterative non-local means denoising ^54^ with *α* = 0.8, patch = 2, and 6 iterations.

### Resolution and CNR calculation

To quantify the resolution the width of a sharp feature transition is usually used, ^36^ but due to the nature of the soft leaf tissue, there are no sharp and well-known features useful for this. Fourier-based correlation method was used to measure the resolution.^46^

### Image analysis and visualisation

All virtual 2D slices were visualized using Fiji. ^55^ Image filtering and 3D visualization were done using Dragonfly. ^56^ All image segmentation and data analysis was done in python.

## Discussion

The data presented above highlight the power of 3D imaging, allowing for non-destructive virtual sectioning through the sample and showing the different tissue types from all angles, deep within the tissue. This type of imaging is widely available and can to some extend even be performed in laboratory setups. For non *in-vivo* experiments where samples can be freeze-dried or chemically fixed, TXM and nano-holotomography are equally good choices for achieving nano-resolution, due to the radiation hardness of freeze-dried samples. Nano-holotomography has a larger field of view than TXM but in many current implementations longer total scan times than TXM and a more complex phase-tomographic reconstruction. Furthermore, TXM imaging is available in lab-based setups and thanks to the simplicity of the tomographic reconstruction it is the obvious choice for initial screening of samples. On the other hand, the focused beam of TXM setups coupled with limited X-ray lens efficiency behind the sample induce radiation damage disrupting tomographic investigation of plants *in-vivo*. In this case we find that live nano-imaging using synchrotron-based nano-holotomography is perhaps the only option today to discover nanoscale aspects in plant physiology in 3D and time. In the lens-less geometry of the nano-holotomography setup, all photons radiated on the sample can be collected by the detector, whereas in TXM ≥ 70% of the photons passing through the sample are lost to inefficient collection in the zone plate. ^57^ Secondly, the high energy X-rays available in the holotomography (29 keV compared to 15 keV in TXM, limited by the efficiency of the Fresnel Zone Plate) means that the absorption cross-section of X-rays in the sample is significantly lower, while the phase signal originating from near field diffraction remains strong. Due to this very high efficiency of radiated photons to collected photons, holotomography is able to image living leaf tissue at down to 50 nm pixel size without causing significant radiation damage to the samples for the time necessary to acquire a handful of tomographic scans at sub 200 nm spatial resolution. Subcellular structures are best visible in the leaf tissue when the pore space between cells is filled with water. This effect occurs due to the high sensitivity of holotomography to phase differences and at the same time the difficulty to reconstruct weak interfaces in the vicinity to strong interfaces in terms of phase shift. In future experiments, using a salt solution for this could potentially reduce cell lysing and allow for imaging of the plant in a less altered state.

## Conclusion

The presented non-invasive approach is instrumental in studying dynamic processes such as water movement, nutrient transport, and structural adaptations, fostering a deeper understanding of plant growth, development, and responses to environmental stimuli without disrupting the natural state of the plant. The mechanisms of loading NPs into the vascular system and their subsequent translocation represent major research gaps within plant science. Future work using these methods will involve tracing the particles as they translocate to the growth zones through the vascular tissue. In the current state, the major challenge lies in choosing a particle designed so that it translocates as an intact particle and, at a determined stage of the translocation process, forms clusters larger than 100 nm. Such studies would benefit greatly by combining with quantitative techniques, such as Small Angle X-ray scattering or X-ray Fluorescence imaging to verify NP size and composition while gaining visual and qualitative insights from nano-tomography. Moreover, with the currently ongoing upgrades of synchrotron sources enabling phase contrast nanoimaging with shorter wavelengths, further reducing the x-ray dose, we foresee an adaptation of the presented approach to a wide range of cases with a focus on studying sub-cellular physiology in 3D.

## Supplementary information

Supplementary information available at: https://doi.org/XX.XXX.XX

## Data availability

All data and videos of data scans used in this publication are available at: https://doi.org/10.11583/DTU.c.7629251. Raw data is available upon request.

## Acknowledgements

We thank Dr. Kentaro Uesugi (SPRING8, BL47XU), Dr. Thorbjørn (MAX IV, DAN-MAX) and Dr. Viktor Nikitin (The Advanced Photon Source) for assisting with experiments and helping with the setup. The European Synchrotron Radiation Facility (ESRF) is greatly acknowledged for provision of beam time (experiment LS3284) using the ID16B beamline. We thank the Danish Agency for Science, Technology, and Innovation for funding the instrument center DanScatt. This research was funded by the Novo Nordisk Foundation (grant no. NNF21OC0066114).

## Declarations

The authors declare no competing interest.

## Footnotes

1 Videos of all scans presented here is available at: https://doi.org/10.11583/DTU.28172201

